# Does Haldane’s rule speciation within mimetic Poison frogs?

**DOI:** 10.1101/2024.03.25.586569

**Authors:** Ugo Lorioux-Chevalier, Mathieu Chouteau, Alexandre-Benoit Roland

## Abstract

To explain how populations with distinct warning signals coexist in close parapatry, we experimentally assessed intrinsic mechanisms acting as reproductive barriers within three *Ranitomeya* poison-frog species. For all species, assortative mating did not occur, nor did a survival disadvantage for inter-population hybrids. However, in *Ranitomeya fantastica*, these hybrids of the male sex are sterile, an outcome predicted by Haldane’s rule. Our results suggest a possible XY sex determinism in *R. fantastica*, and show that this process is associated with an extraordinary diversity, leading to speciation.

**Graphical Abstract:** 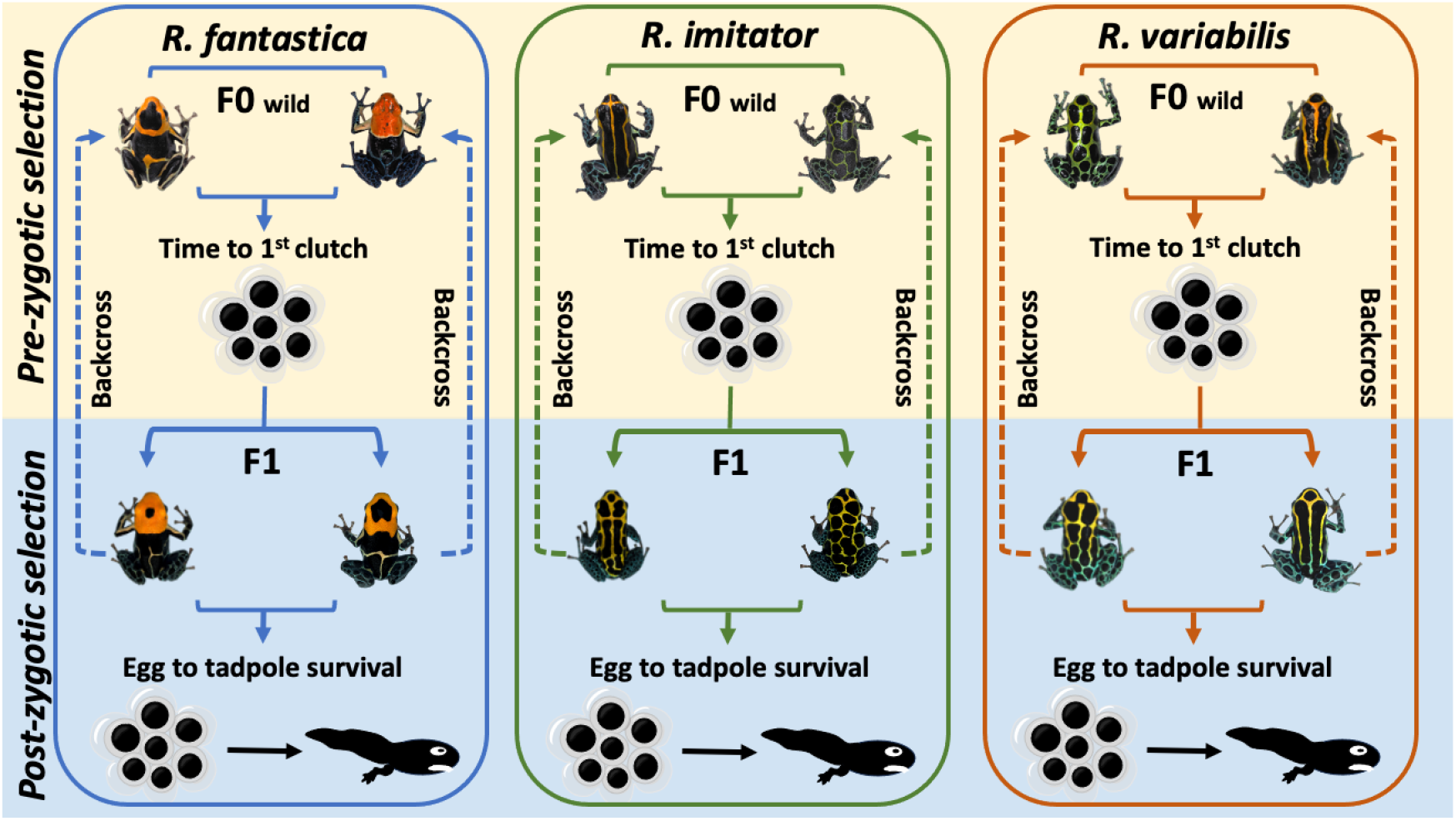

## Introduction

In the evolving landscape of speciation research, the past decades have marked a paradigm shift from geographic isolation to a more nuanced understanding of the mechanisms driving reproductive isolation and diversification amid gene flow ^1^. Divergent selection is recognized as a crucial factor in promoting speciation, supported by both foundational and recent studies ^1–4^. The accumulation of reproductive barriers such as behavioural or gametic isolation, hybrid sterility or unviability is pivotal in adaptive diversification, particularly in interconnected populations ^1,5–8^. In this context, the study of prezygotic and postzygotic barriers sheds light on immediate biological processes and ultimate evolutionary trajectories.

Predation pressure on animal warning signals can create extrinsic prezygotic isolation between populations with distinct colour patterns, as observed in poison frogs ^9^ and butterflies ^10^. Aposematic species face strong natural selection favouring local uniformity, leading to the counter-selection of migrants with exotic signals or hybrid phenotypes. This can drive the evolution of intrinsic reproductive isolation mechanisms such as behavioural selection and genetic incompatibilities, limiting gene flow and recombination between individuals with distinct warning signals ^11^.

In visually driven colour pattern diversification, mate choice leading to assortative mating is hypothesized as a mechanism fostering prezygotic sexual selection ^12^. Indeed, intermediate hybrid phenotypes risk higher mortality due to predator misidentification, favouring the evolution of such reinforcement mechanisms ^13^. On the other hand, postzygotic barriers, such as epistatic incompatibilities leading to hybrid sterility or reduced viability, contribute to reproductive isolation ^8,14,15^, and are crucial in cementing aposematic diversity.

The interplay of prezygotic and postzygotic mechanisms can lead to various scenarios of phenotypic diversification, potentially precipitating speciation events by preventing interbreeding. Understanding these proximate mechanisms is essential for deciphering the ultimate outcomes of species trajectories. Our study explores reproductive isolation mechanisms, among three sympatric clades of aposematic poison frogs in Peru (*Ranitomeya fantastica, R. imitator* and *R. variabilis*). For each species, their impressive diversity of warning signals is geographically structured between adjacent populations ^16,17^, and suggests that intrinsic reproductive barriers might already be in place. This is especially relevant to the *R*.

*fantastica* clade for which it has been suggested they underwent a recent speciation event. Here we unravel the roles of sexual selection and genetic incompatibilities that maintain their amazing local adaptive diversity.

## Results

### Prezygotic barrier

To assess the role of prezygotic barriers, we conducted assays involving pairing similar and dissimilar frog ecotype sequentially, and quantified realized mating events. For all three species tested in the 128 mating experiments, each individual experienced homotypic and heterotypic pairing in random order. Trial success (1) or failure (0) were scored and converted into individual’s preference index (Figure 1). Over the three species and 6 ecotypes tested, only the striped *R. imitator*, seems to have a significant preference for its own phenotype (Supplementary Table 1), while the others do not show any preference. These findings suggest that premating barrier based on colour pattern selection seems to be weak and that frog’s likely do not display assortative mating except for one ecotype of *R. imitator*.

**Figure 1.**
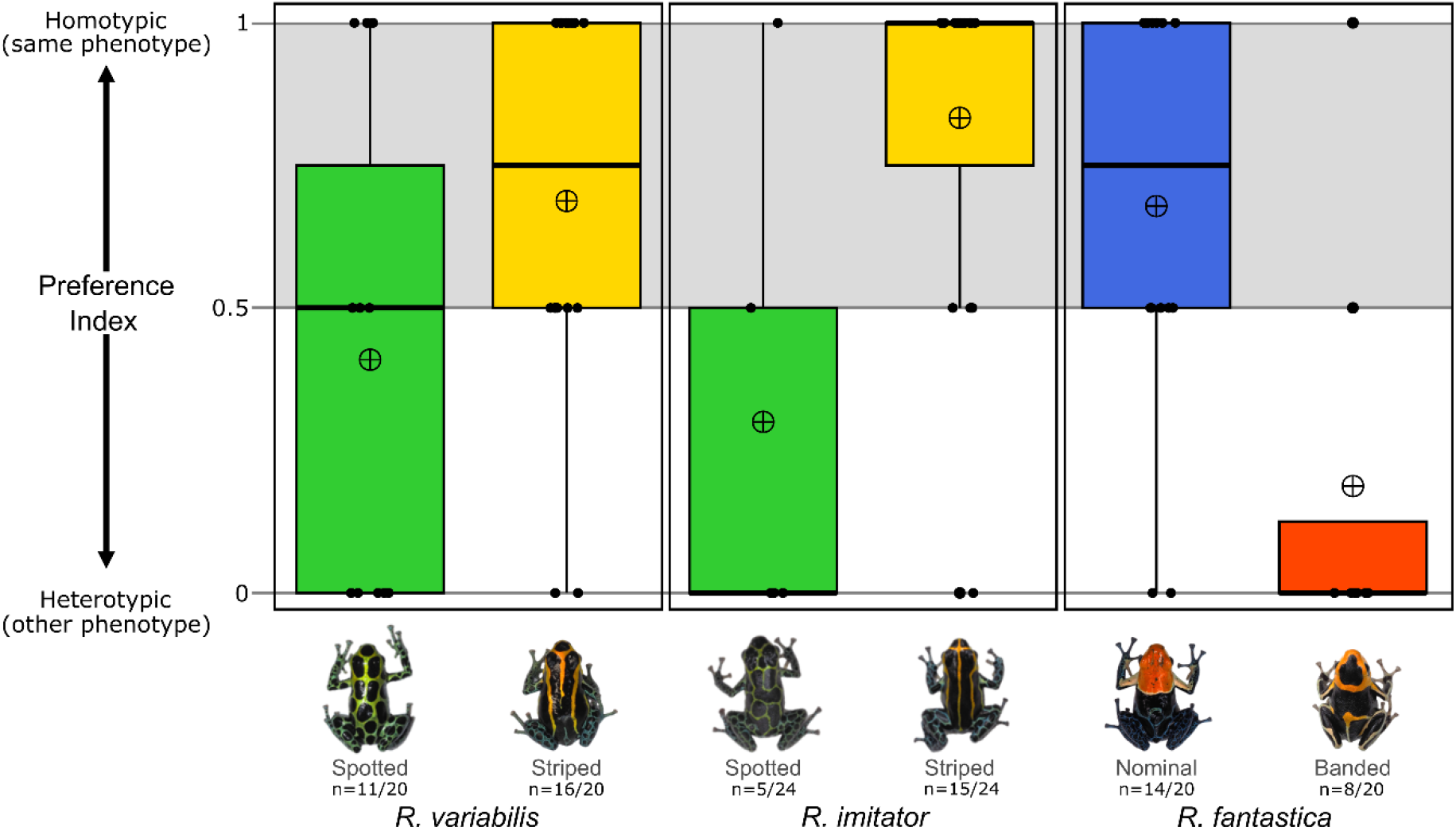
Prezygotic barriers: Individual phenotypic preference. For each animal paired in homotypic and heterotypic reproductive trials for 14 days, success (1) or failure (0) scores was converted into “preference index” (see Methods). Each dot corresponds to one frog and the boxplot presents the mean (cross dot), first and third quartiles, and whiskers extend to ±1.5× the interquartile range. Under each ecotype name is the number of individuals scored over the number tested after removing those that failed both trials.

### Postzygotic barriers

To assess the role of postzygotic isolating barriers, specifically genetic incompatibilities, we conducted homotypic and heterotypic F0 pair crosses. We recorded development time and survival from hatching to metamorphosis. For development time, no significant differences between cross type were observed for *R. variabilis* and *R. fantastica*, but for *R. imitator*, heterotypic tadpoles developed faster (Supplementary Table 2). All F0 pairs produced fertilized clutches and viable F1 offspring (Figure 2). To assess hybrid sterility, we conducted F1_-hybrid_ x F1_-hybrid_ and F1_-hybrid_ x F0_-parental_ crosses. Fertile clutches and viable offspring were produced for every hierarchical cross for *R. variabilis* and *R. imitator*, except for *R. fantastica* (Supplementary Table 3). Notably, all crosses involving F1_-hybrid_ *R. fantastica* males consistently produced clutches with a zero-survival rate (Figure 2). These results indicate that postzygotic epistasis incompatibilities act as a reproductive barrier between the two ecotypes of *R. fantastica*, likely representing distinct taxa. Additionally, the asymmetric sterility of hybrids aligns with Haldane’s rule of speciation suggesting that males could be the heterogametic sex.

**Figure 2.**
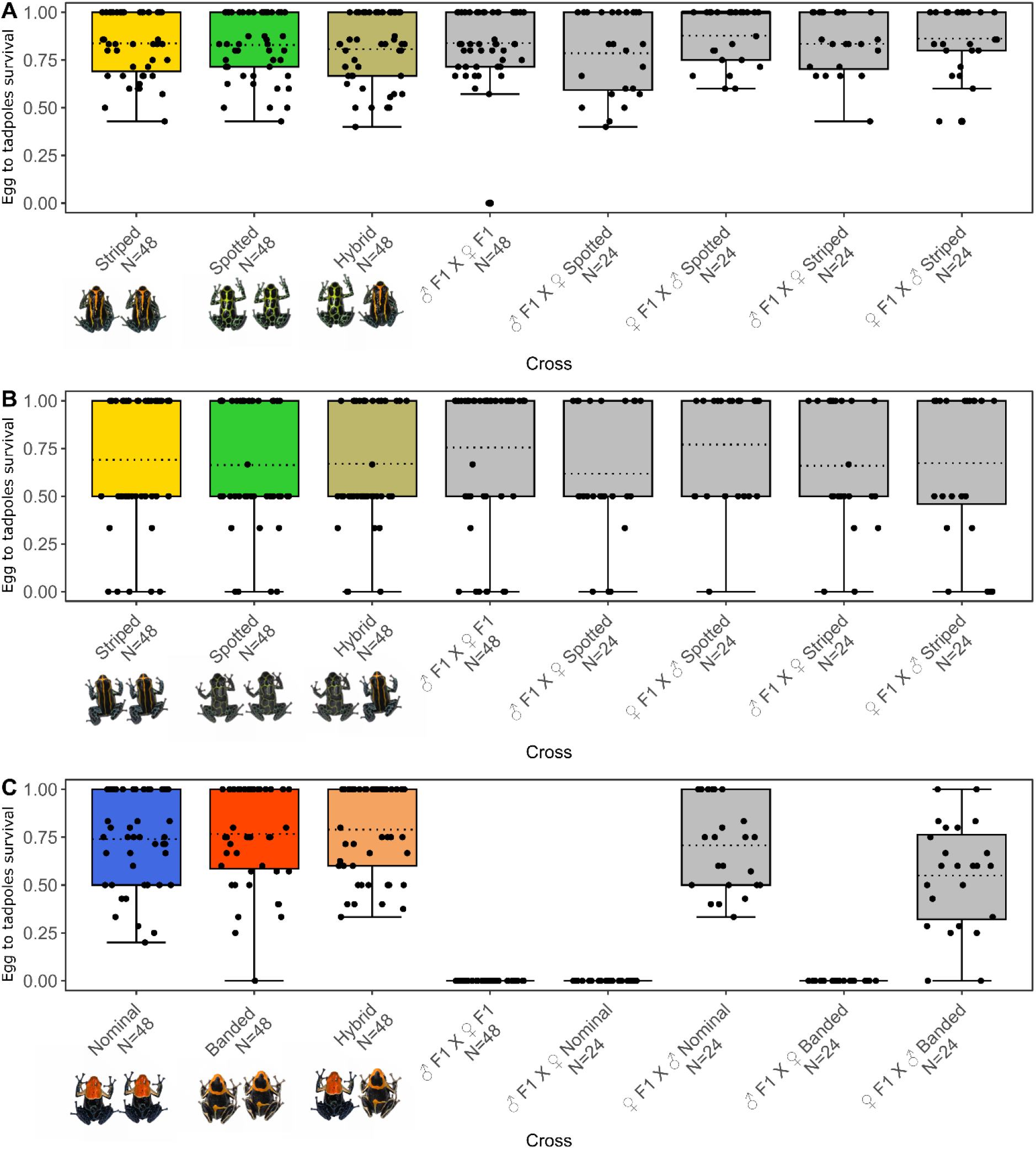
Postzygotic barriers: survival rate of egg to tadpole in F0 and F1 hybrid crosses. Each dot corresponds to one clutch and N represents the number of egg clutches. The boxplot presents the mean in dotted-line, first and third quartiles and whiskers extend to ±1.5× the interquartile range.

## Discussion

Our research delves into the early stages of reproductive isolation and epistasis incompatibilities within the Peruvian *Ranitomeya* system, unraveling both proximate and ultimate mechanisms influencing speciation.

We document the absence of prezygotic barriers through the lack of sexual selection toward similar individuals among *Ranitomeya* species, except for one specific ecotype of *R. imitator* in line with the results of another study ^18^. Instead, we unveil the presence of strong postzygotic mechanisms, suggesting that populations of *Ranitomeya* species are in the early divergence stages ^19^. In *R. fantastica*, the pattern of F1 hybrid male sterility is consistent with Haldane’s rule of speciation, stating that: “When in the F1 offspring of two different animal races one sex is absent, rare, or sterile, that sex is the heterozygous [heterogametic] sex”^20^. Because *R. fantastica* maintains the most diverse mosaic of warning signals compared to *R. variabilis* and *R. imitator* ^16^, these genetic incompatibilities are likely maintaining diversity. These findings indicate that potent epistatic selective pressure against hybrid zygotes, coupled with likely increased predation on the hybrid/exotic warning color pattern, likely contributes to maintaining locally adapted population boundaries by limiting gene flow. This underscores the importance of intrinsic mechanisms in speciation and showcases the diverse nature of reproductive barriers.

One of the two ecotypes of *R. fantastica* of our study was recently categorized within *R. summersi* (our *R. fantastica* “banded” is now *R. summersi* “white banded” ^17^, Supplementary Figure 1). The significant reproductive isolation between our *R. fantastica* populations, consistent with Haldane’s rule, strongly supports their status as distinct species. According to Muell et al., 2022, phenotypic divergence of ecotypes occurs in the same time frame for our three species: *R. fantastica*/*R. summersi (1.78 Mya), R. imitator (0.91 Mya)* and *R. variabilis* (*1.85 Mya)*, but only *R. fantastica* shows genomic signs of speciation. This can likely be explained by the genetic incompatibilities that we unveil, emphasizing the crucial role of intrinsic reproductive barriers in triggering speciation.

Nevertheless, deciphering postzygotic epistasis mechanisms in amphibians poses significant challenges, given that sex chromosome systems often deviate from both Haldane’s rule and the large X-/Z-effect. The widespread application of Haldane’s rule encountered a breach in *Xenopus* frogs, where heterogametic ZW hybrid females, expected to be sterile, remained fertile ^21^. The large X-/Z-effect, which suggests that heterogametic hybrids face genetic imbalance with an X or Z from only one parental species, also faces challenges. This concept typically applies to well-differentiated (heteromorphic) sex chromosomes, as observed in mammals, birds, and certain insects. However, sex chromosomes have frequent recombination in amphibians and behave similarly to autosomes. This unique characteristic possibly leverages the ability of sex chromosomes to switch between male (XY) and female (ZW) heterogametic systems, maintaining a lesser degree of differentiation ^22^. Although resolving the XY/ZW sex determinism is beyond the scope of our study, we present compelling evidence supporting Haldane’s rule in *Ranitomeya*. This insight paves the way for comparative studies, aiming to comprehend the general applicability of these patterns, where amphibian sex chromosome systems challenge conventional ideas in speciation research.

In summary, our research shows that intrinsic prezygotic barriers through sexual selection are not influencing the maintenance of local adaptive diversity despite the fact that positive assortative mating occurs at least in one ecotype of *R. imitator* ^18^. Instead, postzygotic epistasis incompatibilities serve as a reproductive barrier, precipitating speciation in *R. fantastica*. This aligns with Haldane’s rule of speciation, where F1 hybrid males are sterile, suggesting a XY sexual determinism if male prove to be the heterogametic sex.

## Materials and Methods

### Sample collection

Between 2015 and 2022, we gathered a total of 236 wild individuals from three *Ranitomeya* frog species at five locations in Peru (refer to Suppl. Fig. 2 for the localities and names): *R. variabilis* (76 individuals, 2 ecotypes: San-Jose (“Spotted”) and Varadero (“Striped”)), *R. imitator* (84 individuals, 2 ecotypes: San-Jose (“Spotted”) and Varadero Banda (“Striped”), and *R. fantastica* (76 individuals, 2 ecotypes: Micaela (“Nominal”) and Bocatoma (“Banded”)). Each individual was photographed in the lab and assigned a unique ID for identification and tracking within the colony.

### Captive collection and raising conditions

All *Ranitomeya* breeding and crossings were conducted at the Tarapoto Research Center (San Martin, Peru), situated within their natural distribution range. The frogs were housed either in breeding pairs or groups of ten for juveniles, accommodated in 60 x 30 x 30 cm glass terrariums equipped with water drainage, a natural hardscape, and live plants for enrichment. These terrariums were placed in a mesh greenhouse with natural ventilation and a misting system to replicate their native environment. The frogs were fed wingless fruit flies (*Drosophila melanogaster*) and springtails at least three times a week, supplemented with a mix of calcium and vitamins once a week (*Repashy*® *Supercal NoD* and *Supervite)*. Within the breeding enclosures, artificial sites tailored to species-specific needs for oviposition were provided. For instance, *R. fantastica*, which uses covered leaf litter as a deposition site, had 15 cm pipe sections placed horizontally on the ground. *R. imitator*, which uses hollow treeholes or large petioles, had 20 cm pipe sections placed at a 60° angle from the ground to mimic their phytotelma (*Xanthosoma, Heliconia*, etc.). *R. variabilis*, which uses bromeliad phytotelmata, had 5 cm pipe sections sealed on one side placed on the terrarium wall and tilted at a 10° angle, 15 to 20 cm above the ground.

### Prezygotic sexual selection

In *Ranitomeya*, males typically call from exposed perches to attract females during courtship. Realized mating, which reflects the various aspects of mate choice by both sexes leading to a successful mating event, was employed to quantify sexual selection.

Through a straightforward no-choice behavioral assay, mimicking a realistic encountering interaction between a migrant individual with exotic warning signal and a potential mate with the native color-pattern. We recorded the time to he first egg clutch for each pair of frogs over a trial period of 14 days. We conducted a series of homotypic and heterotypic pair crosses for each the three focal species as follows: homotypic (♂_morph-A_ x ♀_morph-A_, n=10-12 biological replicates (pairs) and ♂_morph-B_ x ♀_morph-B_, n=10-12 biological replicates) and heterotypic (♂_morph-A_ x ♀_morph-B_, n=10-12 biological replicates and ♂_morph-B_ x ♀_morph-A_, n=10-12 biological replicates). For each animal paired in homotypic and heterotypic reproductive trials for 14 days, success (1) or failure (0) scores were converted into “preference index” which was calculated as follow:

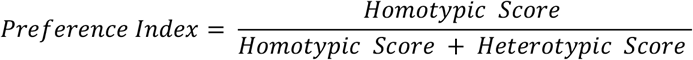

Individuals that failed in both trials were removed to avoid division by zero. Values range from 0 to 1 meaning: 1 being a preference for similar phenotype (positive assortative mating), 0 a preference for dissimilar phenotype (negative assortative mating) and a value of 0.5 shows no preferences. Data normality was assessed using Shapiro test, and we used the nonparametric Wilcoxon signed rank test to compare the median of each ecotype against the hypothetical median (0; 0.5; 1).

### Postzygotic epistasis incompatibilities

To evaluate and quantify the impact of postzygotic epistasis incompatibility on adaptive diversity, we conducted hierarchical crosses for each of the three species. A total of 3513 eggs where followed (1626 for *R. variabilis*, 510 for *R. imitator* and 1377 for *R. fantastica*), and compared between treatments.

Initially, we generated three parental crosses (F0): homotypic (♂_morph-A_ x ♀_morph-A_, n=6 biological replicates; ♀_morph-B_ x ♂_morph-B_, n=6 biological replicates) and heterotypic (F0 Hybrids): ♂_morph-A_ x ♀_morph-B_ (n=3 biological replicates); ♂_morph-B_ x ♀_morph-A_ (n=3 biological replicates). We measured the survival rate (equation below) and development time for each of these crosses, to determine whether heterotypic and homotypic crosses are different (Total clutch number n=432: *R. variabilis* n=144, *R. imitator* n=144, and *R. fantastica* n=144. Total egg number n=1766: *R. variabilis* n=831, *R. imitator* n=265, and *R. fantastica* n=670).

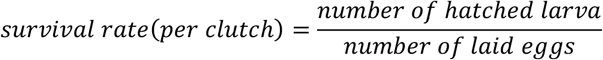

Then we followed a stepwise process for each of the three species, to assess if major epistasis incompatibility results in F1 morph-hybrid adult sterility. Non-related ♂F1_-hybrid_ x ♀F1_-hybrid_ (n=6 biological replicates) crosses were conducted to determine egg development and mortality. To ascertain whether sterility is sex-dependent, we proceeded with ♂F1_-hybrid_ x ♀F0_-morph-A_ (n=3 biological replicates); ♂F0_-morph-A_ x ♀F1_-hybrid_ (n=3 biological replicates); ♂F1_-hybrid_ x ♀F0_-morph-B_ (n=3 biological replicates); ♂F0_-morph-B_ x ♀F1_-hybrid_ (n=3 biological replicates). We measured the survival rate and development time for each of these crosses (Total clutch number n=432: *R. variabilis* n=144, *R. imitator* n=144, and *R. fantastica* n=144. Total egg number n=1747: *R. variabilis* n=795, *R. imitator* n=245, and *R. fantastica* n=707).

Data normality was assessed using Shapiro test, followed by a Kruskal-Wallis and pairwise Wilcoxon test to analyze differences in egg-to-tadpole survival rates between crosses.

## Author contributions

MC conceptualized the study and acquired funding; ULC and MC collected data; ULC and ABR analysed data and prepared the original draft; all authors reviewed and revised the manuscript.

## Acknowledgments

We would like to thank Marco Léon, Henri Delgado, Stephanie Gallusser, Ronald Mori Pezo, Josh Richard, Mario Tuanama from the Peruvian INIBICO NGO for the constant support and logistical help provided during the realization of this project. We also would like to acknowledge the precious comments of Evan Twomey and Kyle Summer on this manuscript. This research was authorized by the Peruvian Servicio Nacional Forestal y De Fauna Silvestre – SERFOR (RDG N° D000442-2021-MIDAGRI-SERFOR-DGGSPFFS and 232-2016-17 SERFOR/DGGSPFFS).

## Data accessibility statement

## Supplementary material

**Supplementary Figure 1.**
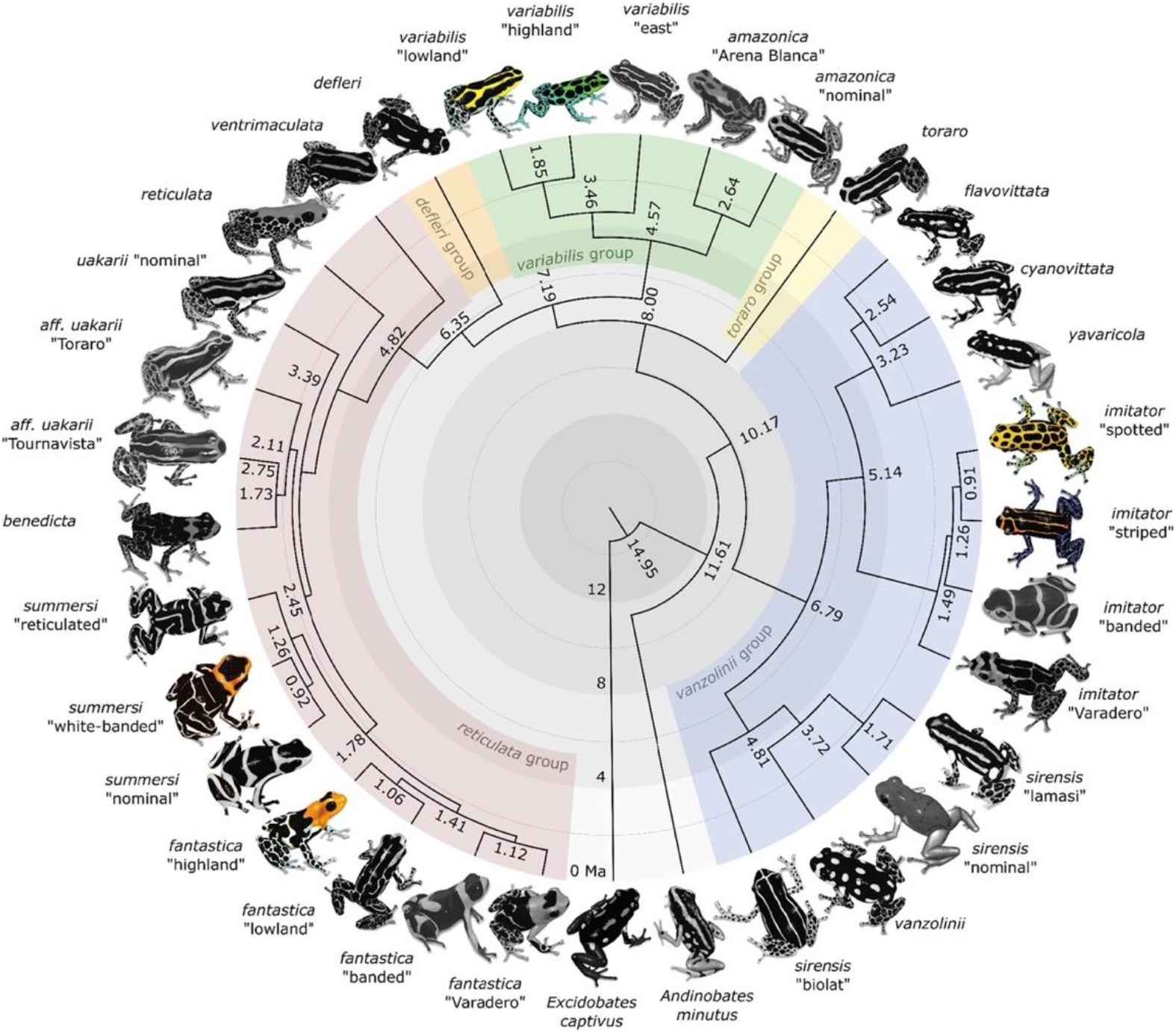
Time-calibrated phylogeny modified from ^17^, highlighting our focal species and ecotypes (colored frogs) within the whole *Ranitomeya* clade (greyed frogs). *Ranitomeya variabilis* “spotted” and “striped” from our study refer to “highland” and “lowland” respectively, from the phylogeny. *Ranitomeya fantastica* “nominal” and “striped” from our study refer to “highland” and *summersi* “white-banded” respectively, from the phylogeny. We used the same denomination for *Ranitomeya imitator*.

**Supplementary Figure 2.**
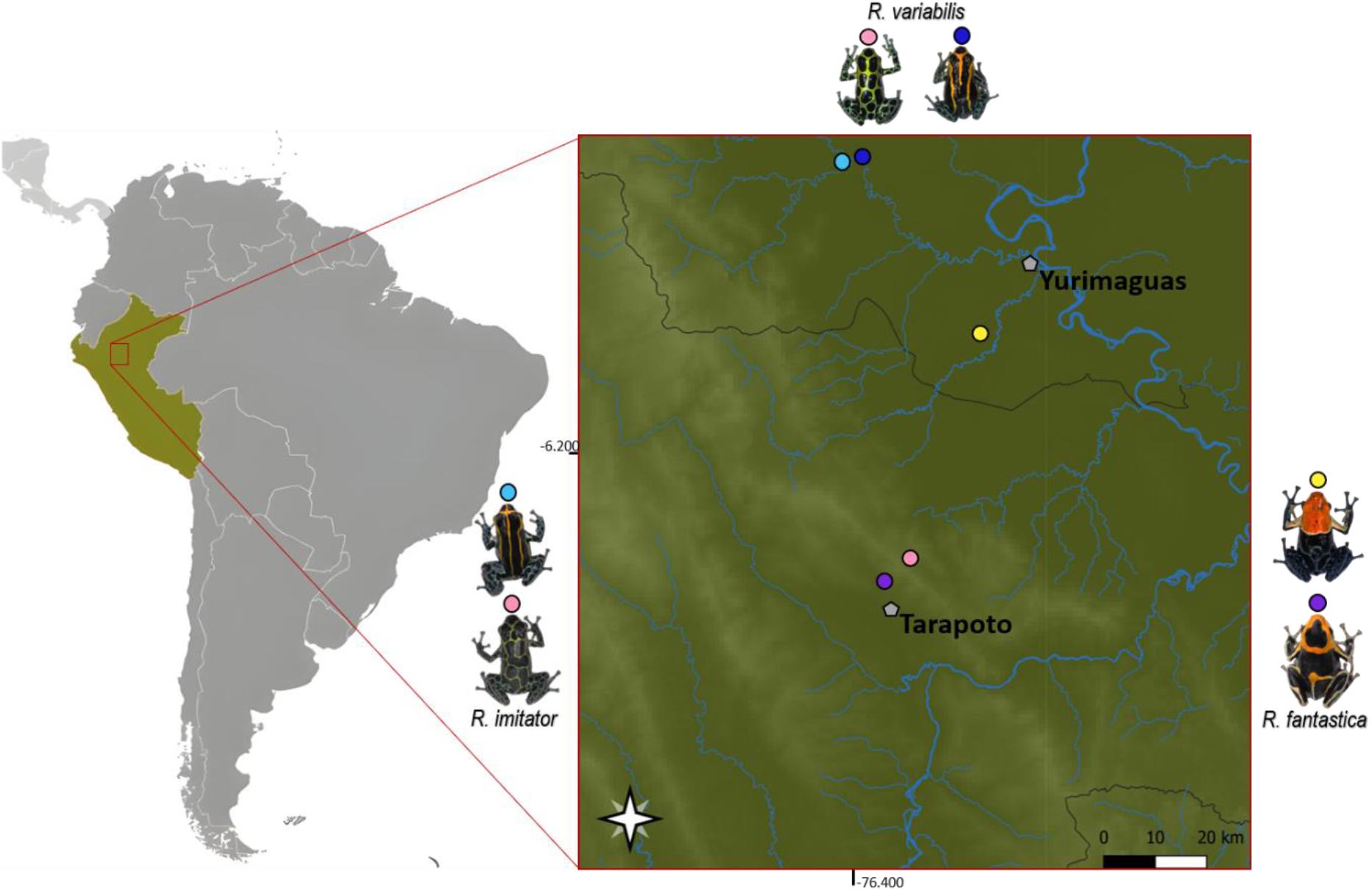
Distribution map of studied species in the region of San Martin and Loreto, Northern Peru, with locality names/ecotypes (Locality name: Purple: Boca-Toma, Yellow: Michaela, Pink: San-Jose, Blue: Varadero, Light-Blue: Varadero-Banda).

**Supplementary Table 1.**
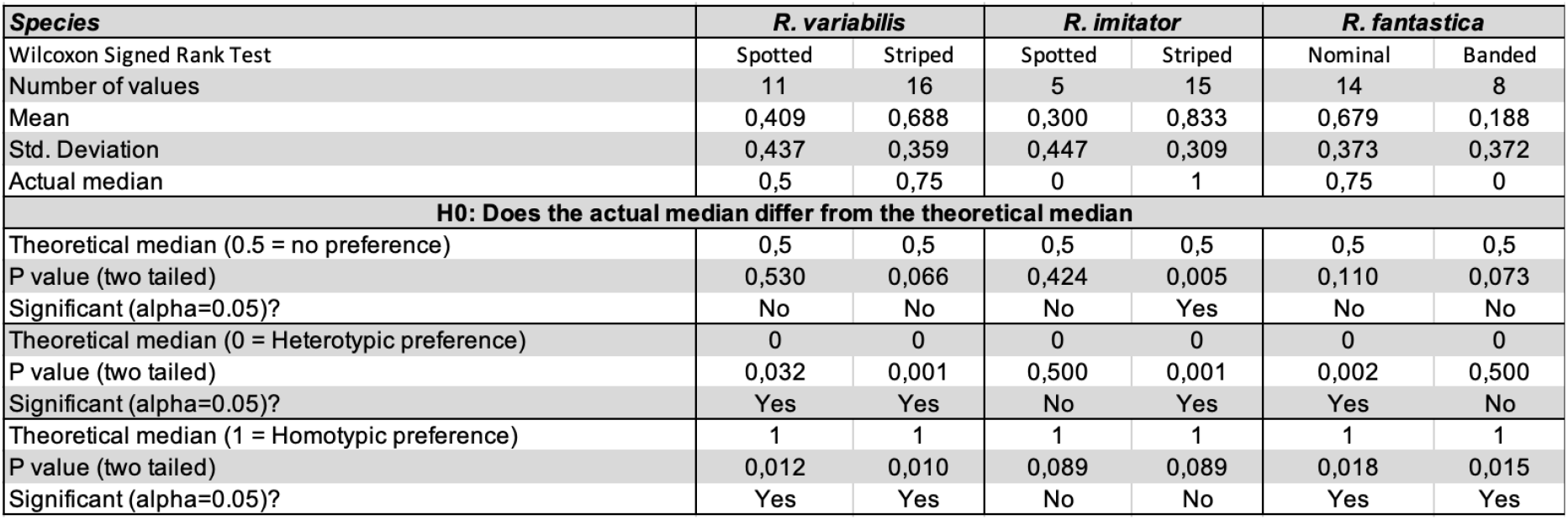
Prezygotic barriers: Individual phenotypic preference. Summary of data and results of the nonparametric Wilcoxon signed rank test.

**Supplementary table 2.**
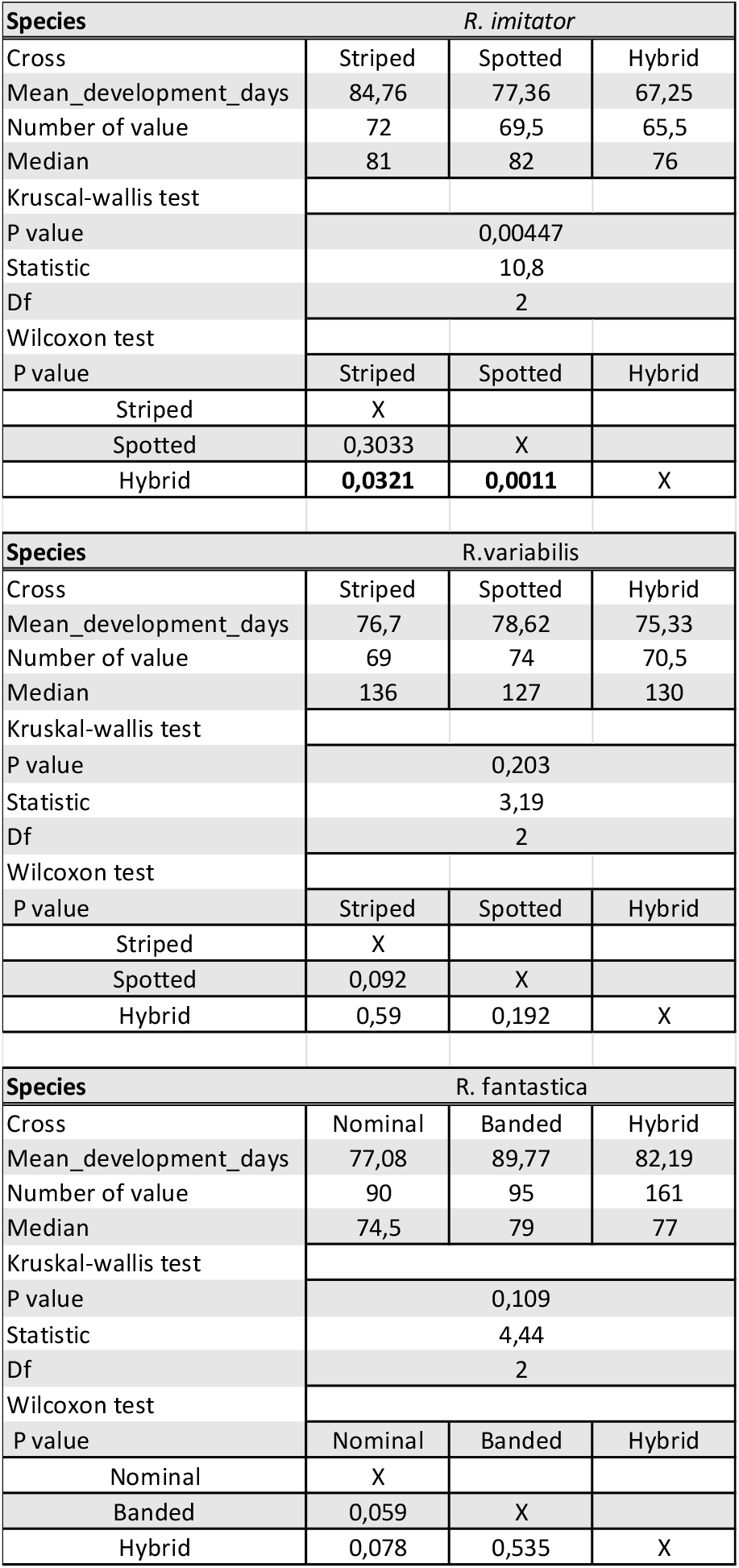
Summary statistics for development time showing the results of the Kruskal-Wallis test and the nonparametric Wilcoxon test.

**Supplementary Table 3.**
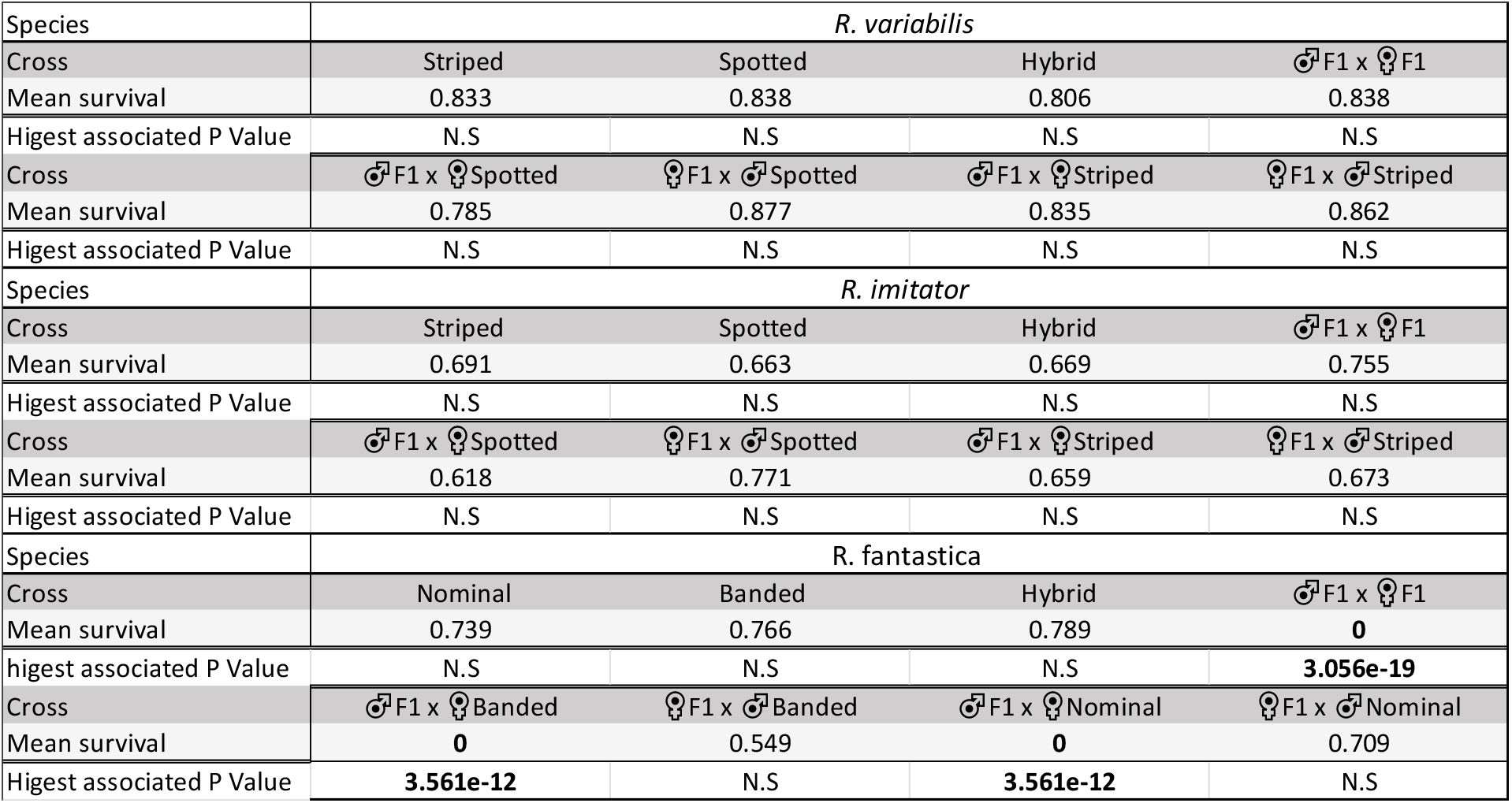
Summary statistics for the survival rate from egg to tadpoles in crossbreed, presenting group means, and their associated p-value. “N.S” for non-significant difference in mean, and bold values when mean was significantly different from controls (in *R. fantastica*: Nominal, Spotted and Hybrid). Only the highest associated p-value is shown.

