## supplementary material for "Does Haldane’s rule speciation within mimetic Poison frogs?"

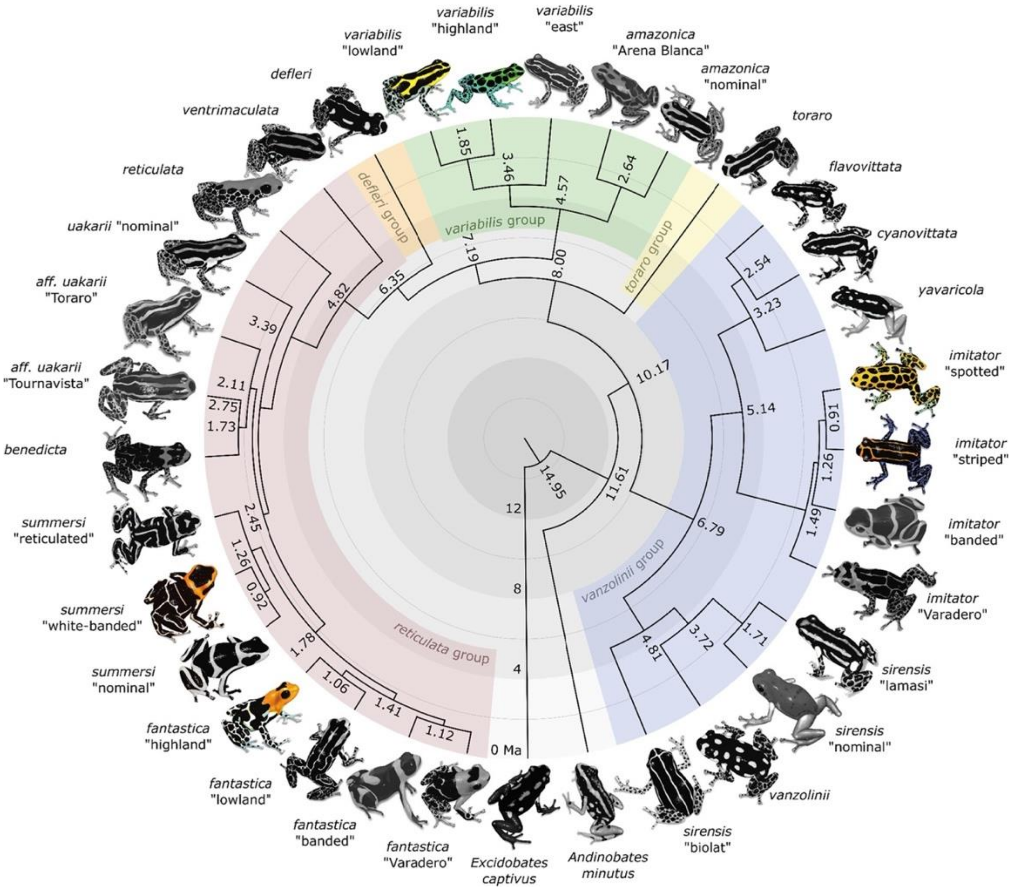


Supplementary Figure 1. Time-calibrated phylogeny modified from [17], highlighting our focal species and ecotypes (colored frogs) within the whole *Ranitomeya* clade (greyed frogs). *Ranitomeya variabilis* "spotted" and "striped" from our study refer to "highland" and "lowland" respectively, from the phylogeny. *Ranitomeya fantastica* "nominal" and "striped" from our study refer to "highland" and *summersi* "white-banded" respectively, from the phylogeny. We used the same denomination for *Ranitomeya imitator*.

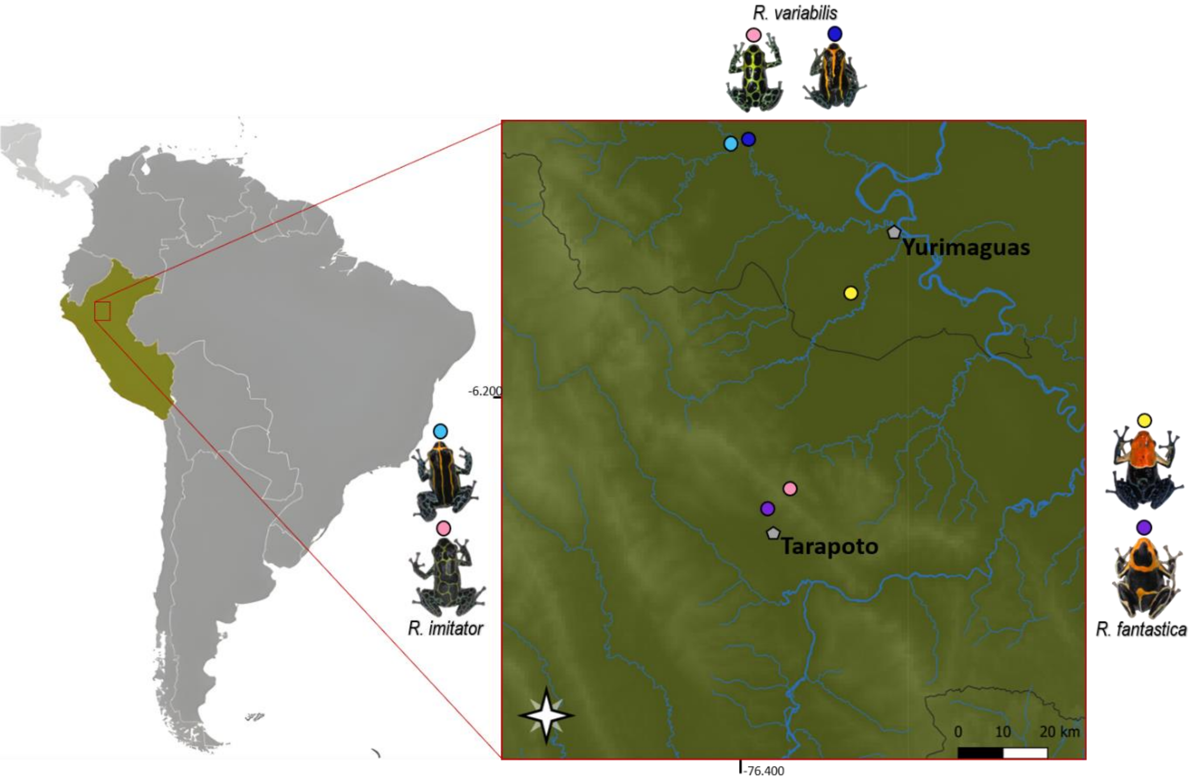


Supplementary Figure 2. Distribution map of studied species in the region of San Martin and Loreto, Northern Peru, with locality names/ecotypes (Locality name: Purple: Boca-Toma, Yellow: Michaela, Pink: San-Jose, Blue: Varadero, Light-Blue: Varadero-Banda).

| ***Species*** | ***R. variabilis*** | | ***R. imitator*** | | ***R. fantastica*** | |
|---|---|---|---|---|---|---|
| Wilcoxon Signed Rank Test | Spotted | Striped | Spotted | Striped | Nominal | Banded |
| Number of values | 11 | 16 | 5 | 15 | 14 | 8 |
| Mean | 0,409 | 0,688 | 0,300 | 0,833 | 0,679 | 0,188 |
| Std. Deviation | 0,437 | 0,359 | 0,447 | 0,309 | 0,373 | 0,372 |
| Actual median | 0,5 | 0,75 | 0 | 1 | 0,75 | 0 |
| **H0: Does the actual median differ from the theoretical median** | | | | | | |
| Theoretical median (0.5 = no preference) | 0,5 | 0,5 | 0,5 | 0,5 | 0,5 | 0,5 |
| P value (two tailed) | 0,530 | 0,066 | 0,424 | 0,005 | 0,110 | 0,073 |
| Significant (alpha=0.05)? | No | No | No | Yes | No | No |
| Theoretical median (0 = Heterotypic preference) | 0 | 0 | 0 | 0 | 0 | 0 |
| P value (two tailed) | 0,032 | 0,001 | 0,500 | 0,001 | 0,002 | 0,500 |
| Significant (alpha=0.05)? | Yes | Yes | No | Yes | Yes | No |
| Theoretical median (1 = Homotypic preference) | 1 | 1 | 1 | 1 | 1 | 1 |
| P value (two tailed) | 0,012 | 0,010 | 0,089 | 0,089 | 0,018 | 0,015 |
| Significant (alpha=0.05)? | Yes | Yes | No | No | Yes | Yes |

Supplementary Table 1. Prezygotic barriers: Individual phenotypic preference. Summary of data and results of the nonparametric Wilcoxon signed rank test.

| **Species** | *R. imitator* | | |
|---|---|---|---|
| Cross | Striped | Spotted | Hybrid |
| Mean_development_days | 84,76 | 77,36 | 67,25 |
| Number of value | 72 | 69,5 | 65,5 |
| Median | 81 | 82 | 76 |
| Kruscal-wallis test | | | |
| P value | 0,00447 | | |
| Statistic | 10,8 | | |
| Df | 2 | | |
| Wilcoxon test | | | |
| P value | Striped | Spotted | Hybrid |
| Striped | X | | |
| Spotted | 0,3033 | X | |
| Hybrid | **0,0321** | **0,0011** | X |

| **Species** | R.variabilis | | |
|---|---|---|---|
| Cross | Striped | Spotted | Hybrid |
| Mean_development_days | 76,7 | 78,62 | 75,33 |
| Number of value | 69 | 74 | 70,5 |
| Median | 136 | 127 | 130 |
| Kruskal-wallis test | | | |
| P value | 0,203 | | |
| Statistic | 3,19 | | |
| Df | 2 | | |
| Wilcoxon test | | | |
| P value | Striped | Spotted | Hybrid |
| Striped | X | | |
| Spotted | 0,092 | X | |
| Hybrid | 0,59 | 0,192 | X |

| **Species** | R. fantastica | | |
|---|---|---|---|
| Cross | Nominal | Banded | Hybrid |
| Mean_development_days | 77,08 | 89,77 | 82,19 |
| Number of value | 90 | 95 | 161 |
| Median | 74,5 | 79 | 77 |
| Kruskal-wallis test | | | |
| P value | 0,109 | | |
| Statistic | 4,44 | | |
| Df | 2 | | |
| Wilcoxon test | | | |
| P value | Nominal | Banded | Hybrid |
| Nominal | X | | |
| Banded | 0,059 | X | |
| Hybrid | 0,078 | 0,535 | X |

Supplementary table 2. Summary statistics for development time showing the results of the Kruskal-Wallis test and the nonparametric Wilcoxon test.

| Species | *R. variabilis* | | | |
|---|---|---|---|---|
| Cross | Striped | Spotted | Hybrid | ♂F1 x ♀F1 |
| Mean survival | 0.833 | 0.838 | 0.806 | 0.838 |
| Higest associated P Value | N.S | N.S | N.S | N.S |
| Cross | ♂F1 x ♀Spotted | ♀F1 x ♂Spotted | ♂F1 x ♀Striped | ♀F1 x ♂Striped |
| Mean survival | 0.785 | 0.877 | 0.835 | 0.862 |
| Higest associated P Value | N.S | N.S | N.S | N.S |
| Species | *R. imitator* | | | |
| Cross | Striped | Spotted | Hybrid | ♂F1 x ♀F1 |
| Mean survival | 0.691 | 0.663 | 0.669 | 0.755 |
| Higest associated P Value | N.S | N.S | N.S | N.S |
| Cross | ♂F1 x ♀Spotted | ♀F1 x ♂Spotted | ♂F1 x ♀Striped | ♀F1 x ♂Striped |
| Mean survival | 0.618 | 0.771 | 0.659 | 0.673 |
| Higest associated P Value | N.S | N.S | N.S | N.S |
| Species | R. fantastica | | | |
| Cross | Nominal | Banded | Hybrid | ♂F1 x ♀F1 |
| Mean survival | 0.739 | 0.766 | 0.789 | **0** |
| higest associated P Value | N.S | N.S | N.S | **3.056e-19** |
| Cross | ♂F1 x ♀Banded | ♀F1 x ♂Banded | ♂F1 x ♀Nominal | ♀F1 x ♂Nominal |
| Mean survival | **0** | 0.549 | **0** | 0.709 |
| Higest associated P Value | **3.561e-12** | N.S | **3.561e-12** | N.S |

Supplementary Table 3. Summary statistics for the survival rate from egg to tadpoles in crossbreed, presenting group means, and their associated p-value. "N.S" for non-significant difference in mean, and bold values when mean was significantly different from controls (in *R. fantastica*: Nominal, Spotted and Hybrid). Only the highest associated p-value is shown.
